# Cell size regulation and proliferation fluctuations in single-cell derived colonies

**DOI:** 10.1101/2022.07.05.498901

**Authors:** César Nieto, César Vargas-García, Juan Manuel Pedraza, Abhyudai Singh

## Abstract

Exponentially growing cells regulate their size by controlling their timing of division. Since two daughter cells are born as a result of this cell splitting, cell size regulation has a direct connection with cell proliferation dynamics. Recent models found more clues about this connection by suggesting that division occurs at a size-dependent rate. In this article, we propose a framework that couples the stochastic transient dynamics of both the cell size and the number of cells in the initial expansion of a single-cell-derived colony. We describe the population from the two most common perspectives. The first is known as *Single Lineage*: where only one cell is followed in each colony, and the second is *Population Snapshots:* where all cells in different colonies are followed. At a low number of cells, we propose a third perspective; *Single Colony*, where one tracks only cells with a common ancestor. We observe how the statistics of these three approaches are different at low numbers and how the *Single Colony* perspective tends to *Population Snapshots* at high numbers. Analyzing colony-to-colony fluctuations in the number of cells, we report an intriguing find: the extent of fluctuations first increases with time and then decreases to approach zero at large numbers of cells. In contrast, in classical size-independent proliferation models, where cell division occurs based on a pure timing mechanism, fluctuations in cell number increase monotonically over time to approach a nonzero value. We systematically study these differences and the convergence speed using different size control strategies.

## I. Introduction

Most cellular properties are subject to random variability (noise) [1]–[3]. Although these fluctuations are traditionally studied in single-cell variables, such as protein levels, stochasticity also affects collective variables involving cell populations such as gene drift [4], [5], host invasion by pathogen cells [6], the survival of small groups to antimicrobial treatment [7], ecology of cell populations [8], size-based interactions [9] and metabolite production [10].

One of the causes of randomness at the population level is the variability in the timing of division [11]. Recent debates have concluded that the rate of division depends on cell size [12]–[23]. Some studies used this size-dependent division rate to partially solve cell size statistics in the cell population [24]–[27] but specifically how this size control affects population dynamics depends on how the population is described.

Two main paradigms are common at modeling population statistics (Fig. 1): the first one is known as the *Single lineage* approach, where the observer tracks only an individual cell of each colony over a large number of colonies (Fig. 1a). Some experiments [22] use the former approach, while others such as cell cytometry [28] and microdroplets [29], use the second perspective: *population snapshots*, which represents the population considering all descendants in a large number of colonies [30], [31] (Fig. 1b).

**Fig. 1.**
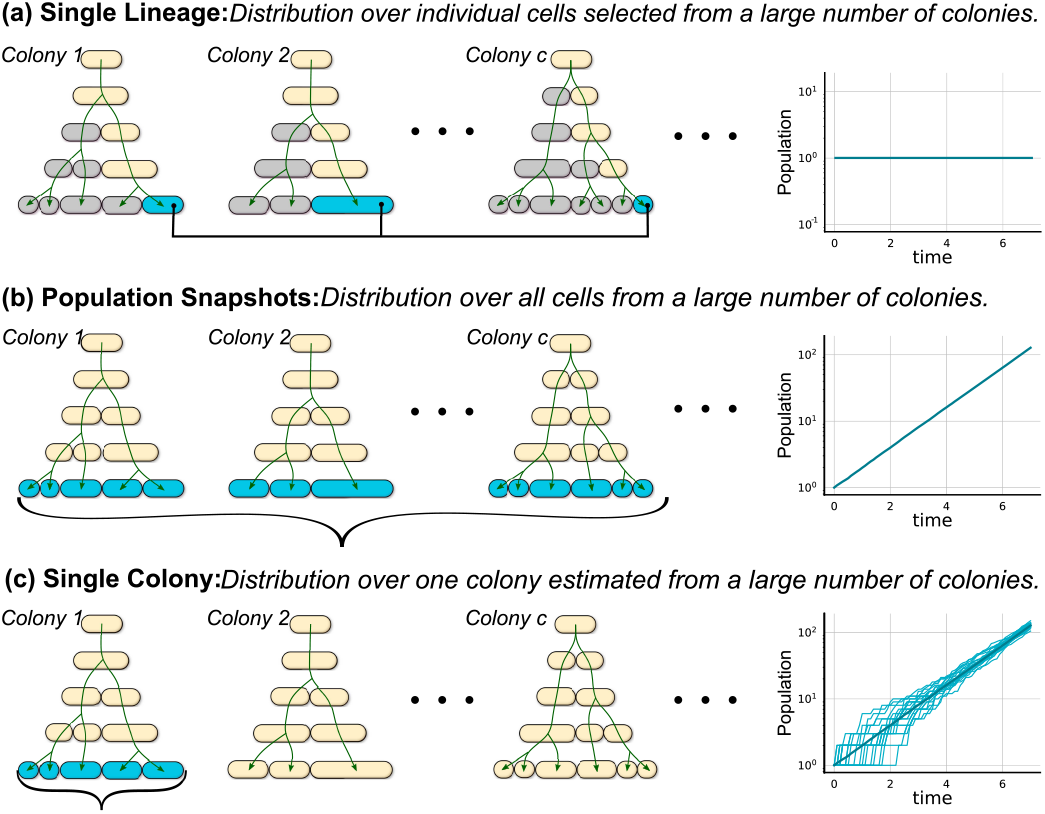
Different approaches to quantify cell size and population count statistics across proliferating colonies. Depending on the perspective, different cells (cyan) are selected to estimate the population statistics.

These perspectives consider that the population has a large number of cells. At this limit, the cell number is taken as a deterministic continuous variable. However, in the initial phase of colony expansion, the number of individuals is low, and its stochastic fluctuations are relatively high. We will discuss how, in this case, it is necessary to propose a third approach: *Single Colony*, where the statistical properties are estimated not over the entire population but across the small number of cells descending from a common ancestor (Fig. 1c).

This article shows methods to calculate the transient dynamics of the cell size distribution for exponentially growing and proliferating cells. We consider these cells to be descendants of a common ancestor with a division rate proportional to size. In Section II, we describe the dynamics of the size distribution using the known Population Balance Equation (PBE), proposing a modification of the finite difference method. We observe how the cell size distribution, despite showing similar moment dynamics, takes different shapes depending on whether we consider random or perfect symmetric partitioning.

In Section III, we model cell proliferation by taking the number of cells as a discrete random variable. We present how, considering cell size dynamics, the *single colony* perspective differs from the other approaches at the beginning of cell proliferation. When the number of individuals is large enough, *single colony* is equivalent to *population snapshots*. Finally, in Section IV, we study the variability of the population between colonies. We compare the size-dependent division model with other models of proliferation based on timer-dependent control. The size dependence reduces the fluctuation in numbers to zero as the population grows. On the other hand, the time-dependent model predicts that these fluctuations do not decrease but instead reach a constant value. [32].

## II. Population Balance Equation

In this section, we will study a numerical solution to the population balance equation (PBE) with some modifications to make it more computationally feasible. The main advantage of this framework is that it estimates not only the moments but also the entire size distribution quickly and efficiently, starting from an arbitrary initial distribution. Recent articles have partially overcome this problem: Some of them found expressions for steady distribution that neglect transient dynamics [26], [30]. Others study the transient dynamics of the distribution moments with limited precision [24]. Finally, further studies can obtain the distribution in exceptional cases and the transient moment dynamics with arbitrary accuracy, but limited to the case of a single lineage [14], [33].

### A. The PBE for growing cells

To introduce the theoretical approach, let us consider the case of an exponentially growing cell without divisions. Let the cell size *s* follow:

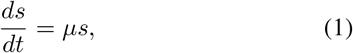

with *µ* being the elongation rate and *t* the time from the beginning of the experiment.

If division is not considered, the associated PBE describing the probability density function (PDF) *ρ*(*s, t*) of the size at any time is given by:

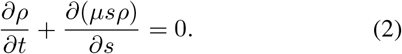

The function *ρ*(*s, t*) can be associated with a probability density function that meets the normalization condition:

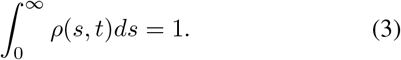

Let the initial condition *ρ*(*s, t* = 0) = *δ*(*s* − *s*_*b*_), that is, the cell starts at an initial size *s*(*t* = 0) = *s*_*b*_. The PBE has the following solution:

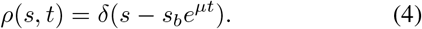

The numerical approach we propose to solve (2) consists of defining an infinitesimal time step Δ*t* that satisfies Δ*t* ≪ *µ*^−1^. Given this Δ*t* and the growth rate *µ*, we can divide the size interval into a variable width lattice that depends on the point *s* as Fig. II-A shows.

**Fig. 2.**
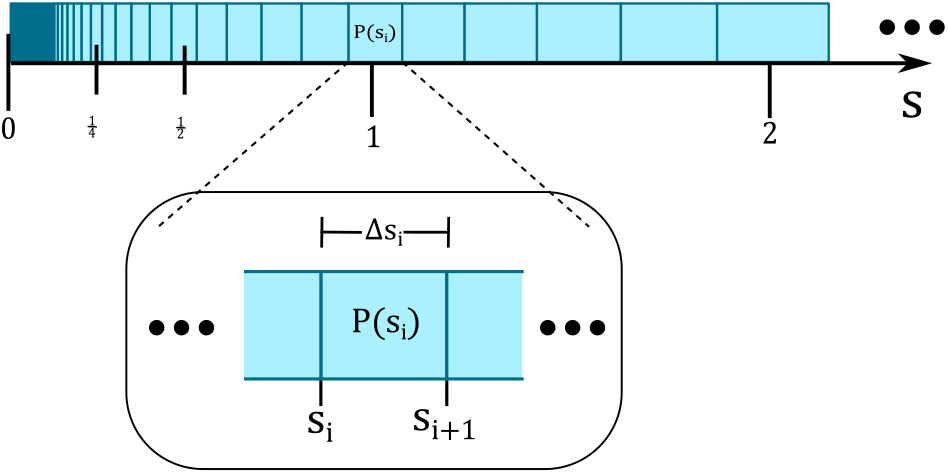
Schematic of the lattice partitioning scheme to solve the dynamics of the PBE equation in exponentially growing cells. The size is divided into a lattice with values *s*_*i*_. The width of the interval (*s*_*i*_, *s*_*i*+1_) = (*s*_*i*_, *s*_*i*_ + Δ*s*_*i*_) is proportional to *s*_*i*_ by the relationship Δ*s*_*i*_ = *s*_*i*_*µ*Δ*t* with Δ*t* being an infinitesimal time step used in integration as explained in (17).

After discretization of (1), the size *s*_*i*+1_ with *i* ∈{0, 1, …, *I*} can be defined from the size *s*_*i*_ using:

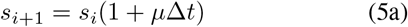

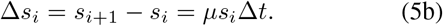

If division is not considered and using (5a), the expression (2) can be written as:

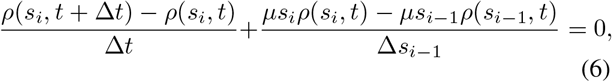

which, after multiplying on both sides by Δ*t*Δ*s*_*i*_, it becomes:

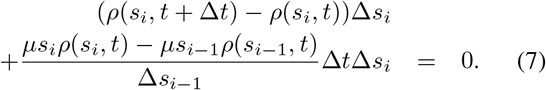

After using the properties 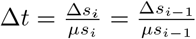 and Δ*s*_*i*_ = Δ*s*_*i*−1_(1+*µ*Δ*t*) derived from (5b) and (5a), respectively, we define the probability of finding cells of size in the interval (*s*_*i*_, *s*_*i*+1_).

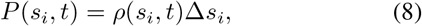

(7) can now be expressed as:

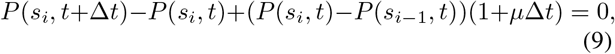

which, after the condition *µ*Δ*t* ≪ 1 simplifies into:

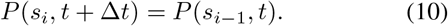

This expression, together with the initial condition 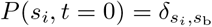 with *δ*_*i,j*_ being the Kronecker delta, is equivalent to (4).

### B. The PBE for growing and dividing cells

Here, we will present how to modify the solution for PBE, including division. First, consider exponential growth as (1) and the division rate *h*(*s*) as proportional to the size as *ks*. Under these assumptions, the associated PBE that describes the number density function *n*(*s, t*) follows:

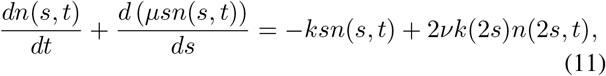

where, as explained in [24], the parameter ν defines the perspective used to approach population statistics. ν = 1, for instance, corresponds to the perspective of a single lineage, while ν = 2 is used to describe population snapshots.

In this case, the PBE describes the evolution of the function *n*(*s, t*) instead of a PDF *ρ*(*s, t*). This is because *n*(*s, t*) satisfies a different normalization condition:

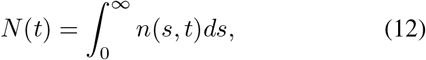

with *N*(*t*) usually interpreted as the total of cells in the population that defines the PDF as [24]:

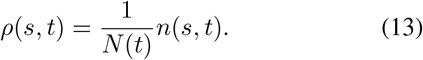

The term to the right of (11) after performing similar procedures to obtain (7) can be discretized as:

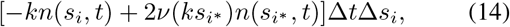

where we define the subindex *i*\* as *s*_*i*\*_ ≈ 2*s*_*i*_. To estimate *i*\*, we can use the result (5a) to obtain

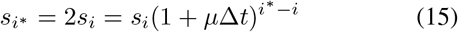

that can be used to solve *i*\* as:

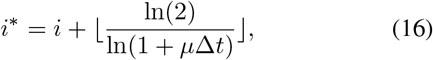

with ⌊*x*⌋ being the highest integer less than or equal to *x*.

With formula *N*(*s*_*i*_) = *n*(*s*_*i*_)Δ*s*_*i*_, the definition of 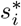 in (15), and noting that 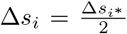, the expression (11) can be written as:

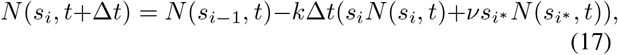

where the function *N*(*s*_*i*_, *t*) defines the number of individuals expected in the size interval (*s*_*i*_, *s*_*i*+1_) at a time *t*.

After considering an arbitrary starting distribution *N*(*s, t* = 0), applying the time evolution given by (17), as Fig. 3 shows, one can obtain the size distribution *N*(*s*_*i*_, *t*) at any time later in the future.

**Fig. 3.**
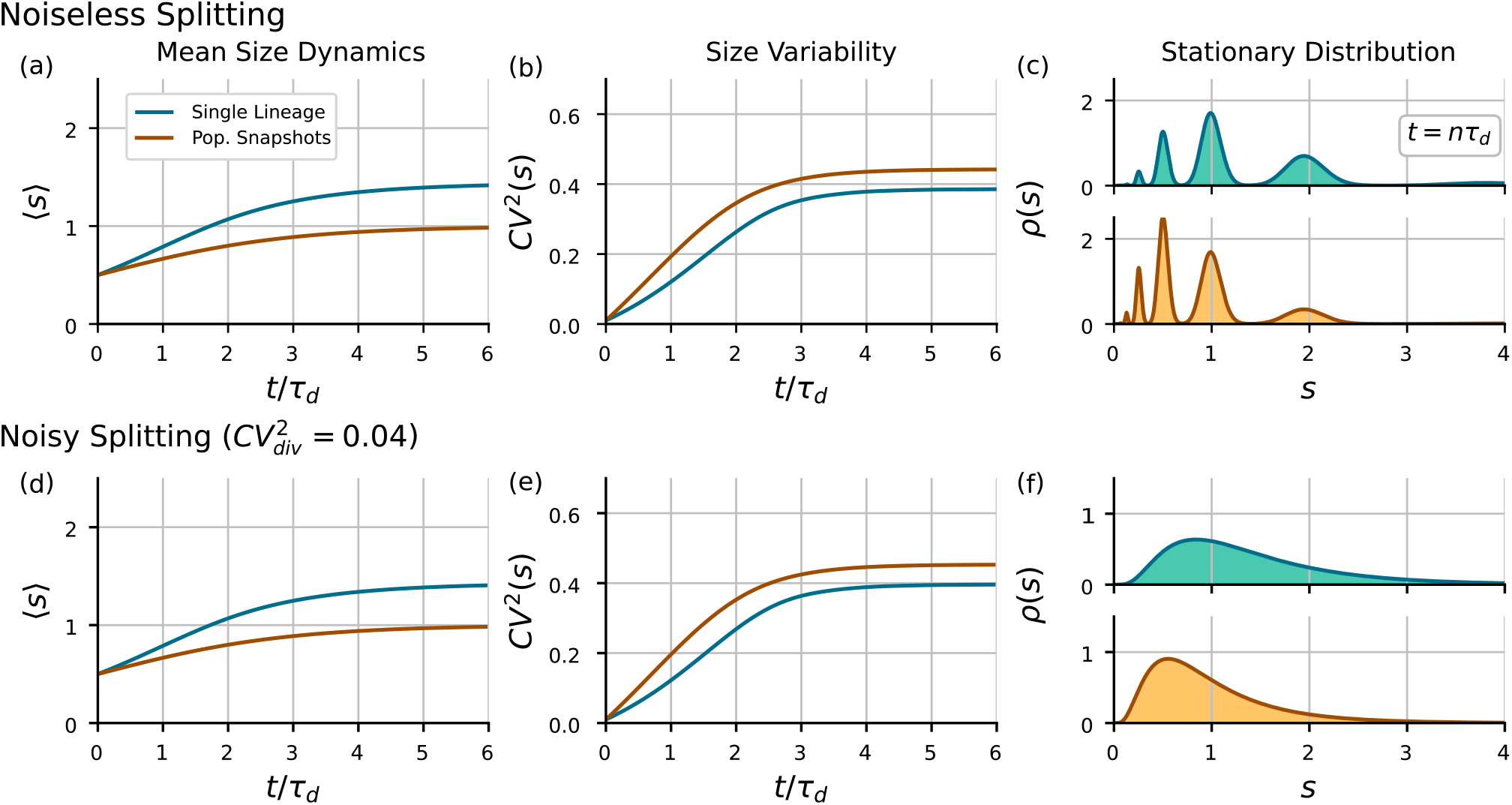
Differences between the distribution dynamics depending on the perspective used for doing the size statistics. *Top:* (a) Mean size dynamics, (b) size variability measured as *CV* ^2^(*s*), and (c) the size distribution observed for time equal to a multiple *n* of the doubling time *τ*_*d*_. *Bottom:* Stochastic dynamics of the size distribution, including noise in partitioning 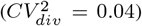. (d) Mean size dynamics, (e) size variability measured as *CV* ^2^(*s*), and (f) the size distribution observed for time equal to a multiple of the doubling time *τ*_*d*_. The single lineage perspective is shown in blue, whereas the population snapshots case is shown in brown. (Parameters: *k* = *µ* = ln(2), Δ*t* = 0.0025, *s*_0_ = 2^−6^, *I* = 8000, ⟨*s*⟩|_*t*=0_ = 0.5, *CV* ^2^(*s*)|_*t*=0_ = 0.01, *i*\**i* = 400)

The mean population *N*(*t*) over time, the mean size ⟨*s*⟩, and the variance var(*s*) are given by the following:

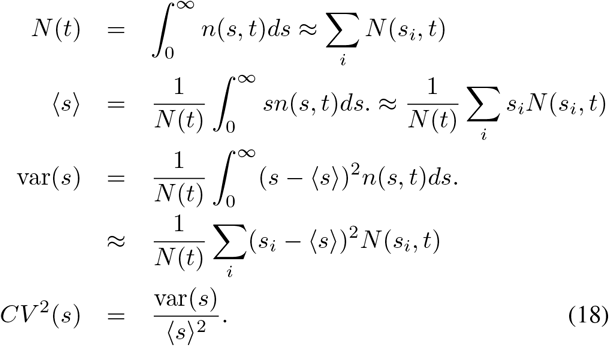

As we explained in previous studies [14], we normalize the time by the doubling time such as *τ*_*d*_ = 1, that is, *µ* = ln(2) and the size by the mean added size before division 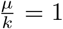 so, we chose *k* = *µ* = ln(2). Thus, all sizes are measured in units of *µ/k*.

In Fig. 3, we present the moment dynamics of the size distribution from an initial size *s*_*b*_ := *s*(*t* = 0) following a gamma distribution with parameters ⟨*s*_*b*_⟩ = 0.5 and *CV* ^2^(*s*_*b*_) = 0.01. As previously found, the steady mean size for population snapshots is lower than this value for the single lineage (see Fig. 3a) [24], [34]. This effect is expected since, from the snapshots perspective, the fastest proliferating cells are smaller and more abundant. The size variability *CV* ^2^(*s*) reaches similar values for both perspectives with higher fluctuations for the population snapshots perspective (Fig. 3b). Unlike other articles [24], in this approach, we also present the limit distribution (Fig. 3c). Although it has a steady mean ⟨*s*⟩ and steady *CV* ^2^(*s*), the distribution presents a limit cycle, as observed in previous studies [14], [25]. The complete time dynamics can be found in Supplementary videos 1 and 2 for the single lineage and population snapshots, respectively.

### C. Time evolution considering noise in splitting

Cells do not split perfectly in half, but with some random fluctuations. As we suggested before [14], this noise in the splitting position may break the limit cycle leading to a stationary distribution. To observe how this stochasticity affects the transient dynamics of the cell size distribution, we propose the associated PBE equation, including partition noise [26]:

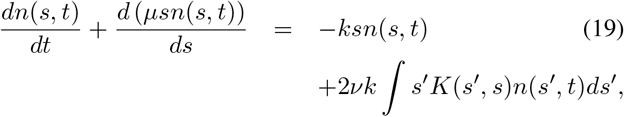

where *K*(*s*′, *s*) is a kernel function. We assume that after division, the size is not divided by one-half but multiplied by a constant *b* following:

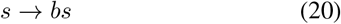

where ⟨*b*⟩= 0.5. We can assume, for example, that *b* follows a Beta distribution with a given *CV*. To solve (19) we calculate the value of the beta distribution over the interval (0.25, 1) to obtain the following:

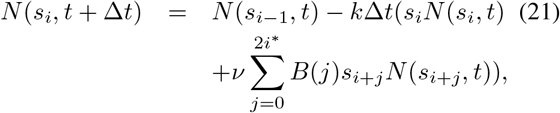

with *B*(0) = 0.25*µ*Δt*β*(0.25) and *B*(2*i*\*) = *µ*Δ*tβ*(1) with *β*(*x*) the beta distribution value at point *x*.

The resulting moment dynamics using the algorithm (21) is presented in Fig. 3d and 3e. Although we have relatively high noise in the splitting position (20%), we can observe that the dynamics does not change appreciably. However, the size distribution reaches a stationary shape (Fig. 3f), which looks completely different from the case of noiseless splitting. Supplementary videos 3 and 4 present these cell distribution dynamics for single lineage and population snapshots, respectively.

### D. Comparison with data sets

In previous sections, we explain how to obtain the size distribution for different perspectives of population description. A direct comparison between these trends and the data is not straightforward. In this article, we contrast the moments in the system (18) with those derived from data (simulations) by estimating the moment ⟨*s*^*α*^⟩ from *C* colonies, each with *Z*_*c*_ cells, with cell size *s*_*z,c*_:

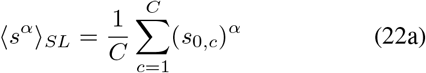

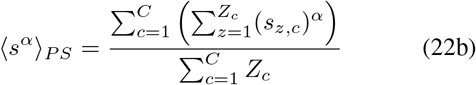

with *SL*, referring to single lineage perspective, and *PS* referring to Population Snapshots. These formulae were also used to compare the solutions of numerical methods and stochastic simulation algorithms [14], [35].

This section presented an algorithm for efficient computation of the size distribution dynamics for exponentially growing and dividing cells. This framework considers any arbitrary starting distribution and includes noise in the splitting position. Calculating the entire size distribution over time allows us to better understand the transient dynamics of cell proliferation. However, this approach does not allow for distinguishing between cells that descend from a common ancestor, since PBE includes all cells in many colonies. We propose a new framework to overcome this issue, as seen in the next section.

## III. Single Colony Statistics

To model a growing population of cells that divide exactly in half, consider the state 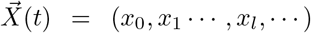 defined by the number of cells with *l* divisions after a time (*t*) as presented in Fig. 4. For example, the vector 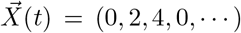 defines the state of six cells since Σ_*i*_ *x*_*i*_ = 6. Two of these cells have divided once (*x*_1_ := 2), while the other four cells have divided twice (*x*_2_ := 4). There are no cells without divisions (*x*_0_ := 0) and there are no cells with more than two divisions (*x*_*l*_ := 0, *l* > 2).

**Fig. 4.**
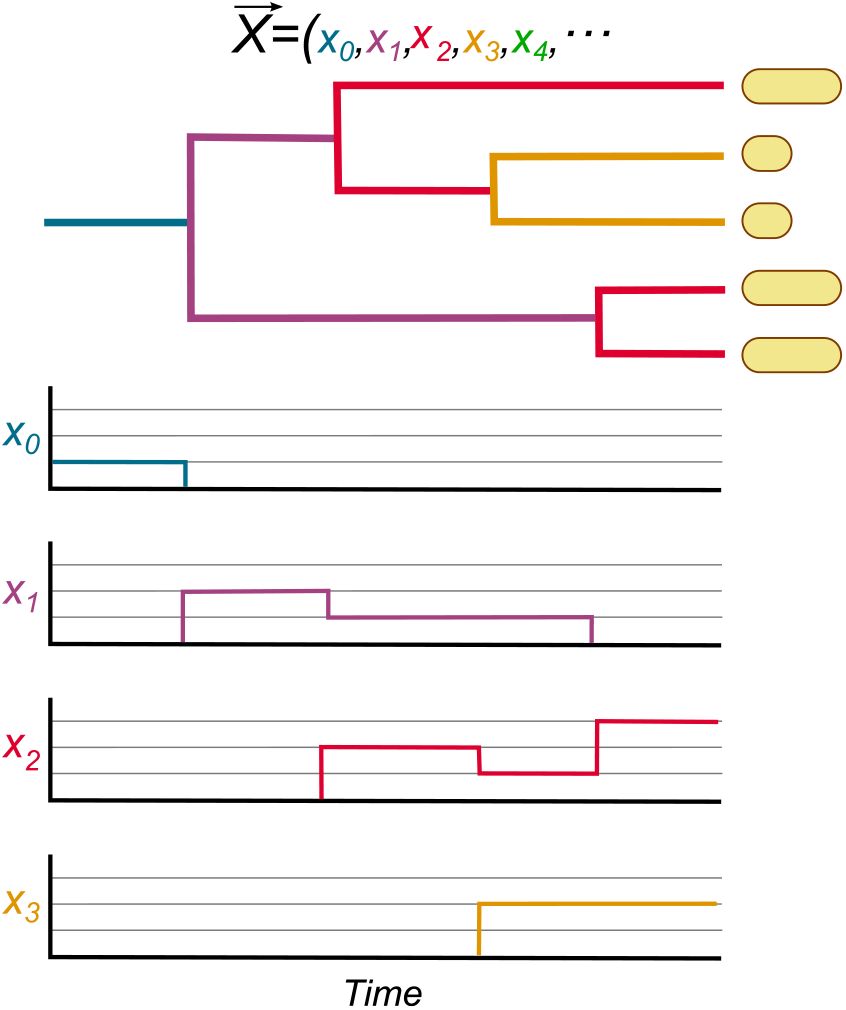
Definition of a state 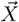 describing the population of cell. In this diagram, we show how 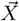 changes in a proliferating population. *Blue*-. Number of cells without division xq. *Purple:* Cells with one division x_0_. Red: Cells with two divisions x_2_. *Yellow:* Cells with three divisions x_3_. Green: Cells with four divisions. At the end of the proliferation process, the colony is described by the state 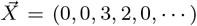 and the total number of cells is Σ_*i*_ *x*_*i*_ = 5.

As explained in previous articles [14], [25], consider cells descending from a single ancestor that had a cell size *s*_*b*_ at time *t* = 0. If they are growing exponentially at rate *µ*, the size of a cell after perfectly symmetric divisions *l* is:

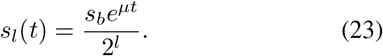

Now, consider the division event for cells of size *s*_*l*_. If there are *x*_*l*_ cells of this size, each of them dividing at rate *ks*_*l*_, the event consisting of one of these *x*_*l*_ cells divides, indistinguishably, at rate *x*_*l*_(*ks*_*l*_). After division, the number *x*_*l*_ decreases by one and *x*_*l*+1_ increases by two since two new cells are born. This division is described by the following transition:

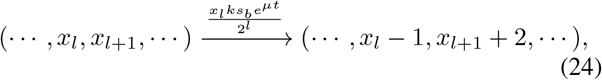

and other transitions to vectors different from (24) are forbidden.

Defining the state 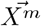 as a population state prior to division of one of the cells that has divisions *m*, we have the following.

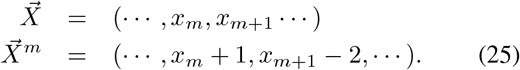

The master equation describing the dynamics is given by:

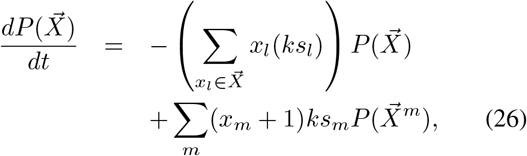

where *s*_*l*_ follows (23).

As an example of (26), we can show the master equation for the first three states:

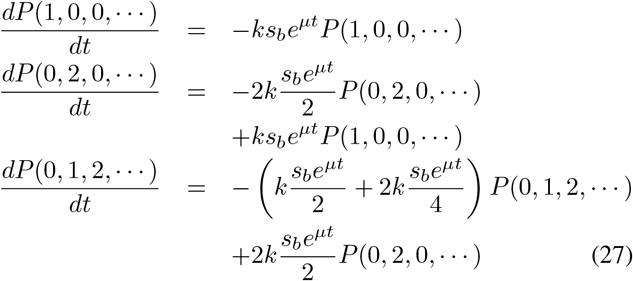

Numerical methods based on the finite state projection algorithm [36] were used to obtain a computation of 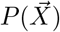 over time. This algorithm truncates the number of states, in principle infinite, to a given number of finite numbers and integrates the master equation (26). In Fig. 5, we show the dynamics of cell size moments with projection over the first seven generations. Given that the number of possible states that truncate the number of divisions up to *L* increases 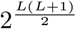, it was not possible to increase the accuracy of the algorithm for times greater than 5*τ*_*d*_.

**Fig. 5.**
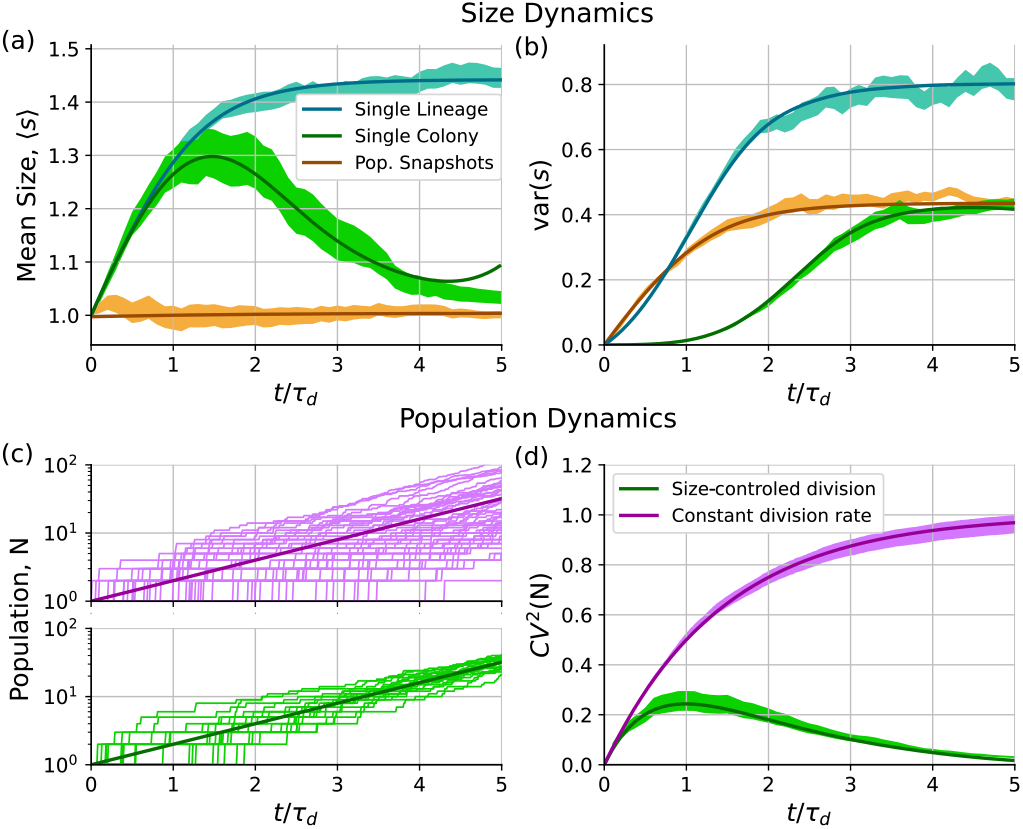
Moments dynamics of both the cell size distribution and the population numbers for the different population perspectives. (a) Dynamics of the mean size using (28) for a single colony perspective (green) compared to the obtained using (17) for a single lineage (blue) and population snapshots (brown). The variance of the size distribution (b) is also compared in the three perspectives. The simulations used the formula (22) and the bootstrapping methods to estimate the confidence intervals of the simulation results. (c) Examples of population number trajectories for a constant division rate (purple) and for a division rate proportional to the size (green). The thick lines correspond to the mean population. (d) Dynamics of the variability of the population number measured using *CV* ^2^(*N*). The lines represent the numerical results (30) and the colored regions are 95% confidence intervals of the results of stochastic simulations (31. (Parameters: *k* = *µ* = ln(2), ⟨*s*⟩|_*t*=0_ = *s*_*b*_ = 1, *CV* ^2^(*s*)|_*t*=0_ = 0, ⟨*N*⟩_*t*=0_ = 1, *CV* ^2^(*N*)|_*t*=0_ = 0).

### A. Size distribution moment dynamics

Once 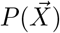 is estimated, the moments of the size distribution ⟨*s*^*α*^⟩_*SC*_, where the subscript *SC* stands for single colony, can be calculated as follows:

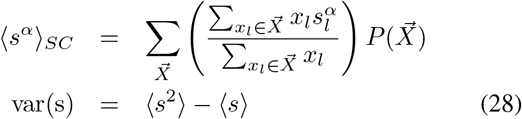

These moments can be compared with the empirical value estimated to have *C* colonies, each with *Z*_*c*_ cells of size *s*_*z,c*_:

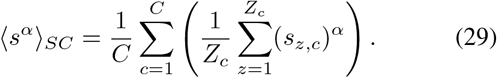

Fig. 5 shows how the dynamics of the moments of the size distribution differ depending on which of the three perspectives we choose. In addition to the moment dynamics already observed using the PBE and equation (18), in Fig. 5a, we observe how the mean size ⟨*s*⟩ from the perspective of a single colony follows a dynamics similar to the single lineage, and after four division times, it is equivalent to population snapshots. Size variability, measured as the variance of the size distribution (Fig. 5b), shows how, from the perspective of a single colony using (28), this var(*s*) remains null-valued during the first doubling time. This effect occurs because the cells are perfectly split in half and both daughter cells have the same size after the first division.

In this section, we theoretically observe a model to approach the size distributions across single-colony descriptions. This model is a generalization of previous approaches [25], where the number of cells with a given number of divisions defines the state of the system. The numerical solution of the associated master equation allowed us to estimate the moment dynamics of the size in the three perspectives of population description. In the next section, we will study the stochastic dynamics of the number of individuals in the colony. We will also compare the results of this size-dependent division with classical models, where the division occurs at a size-independent rate.

## IV. Dynamics of the variability on population number

The variability of the population number can be estimated from the moments of the population number *N*:

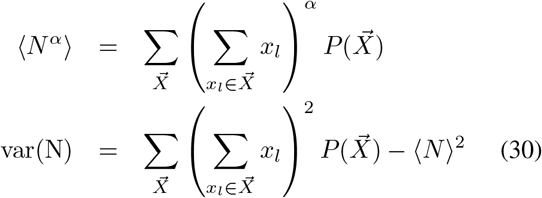

These moments can be compared with the estimates from data sets with considerations similar to those used in (22):

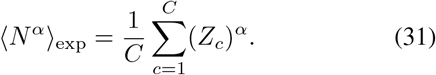

In Fig. 5c, we show some trajectories of population numbers for two systems. One of them (purple) consists of cells that divide at a constant rate, which is a widely known model in proliferation dynamics [37], [38]. As a main result, Fig. 5d shows how the size-controlled division model presents an asymptotic decay to zero in *CV* ^2^(*N*). On the other hand, in the model with a constant rate of division, this variability increases to 1.

### A. Population fluctuations in cell size-controled vs timercontroled proliferation

To better understand the models of division timing, let us compare two simple mechanisms of cell division. If the division is size-controled, the splitting event is decided based on the cell size. On the other hand, the division is timercontroled if the cells consider the timer *τ*. This timer is defined as the time spent since the most recent division.

The division rate is modeled by the function *h*(*s, τ*) as recently proposed [26], [39]. Therefore, the probability of dividing during the interval (0, *τ*) is given by:

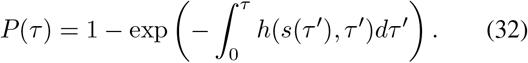

In the particular case where the size *s* is used as a reference to divide, the division rate can be expressed as follows.

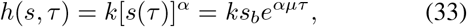

with 0 < *α* < ∞, an exponent defining the strength of the control [18], [26]. In this case, the distribution of division times depends on the size at the beginning of the cycle *s*_*b*_ [14]:

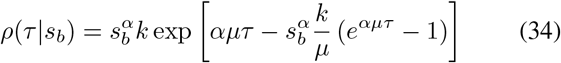

In our simulations, we generate random times *τ* from the PDF (34). During the time interval (0, *τ*), cells grow exponentially following (1) and split in half with partition noise (*CV* ^2^(*b*) = 0.005). Fig. 6b shows how, as *α* increases, *CV* ^2^(*N*) converges faster to zero. The special case *α* → 0 corresponds to the division at a constant rate.

**Fig. 6.**
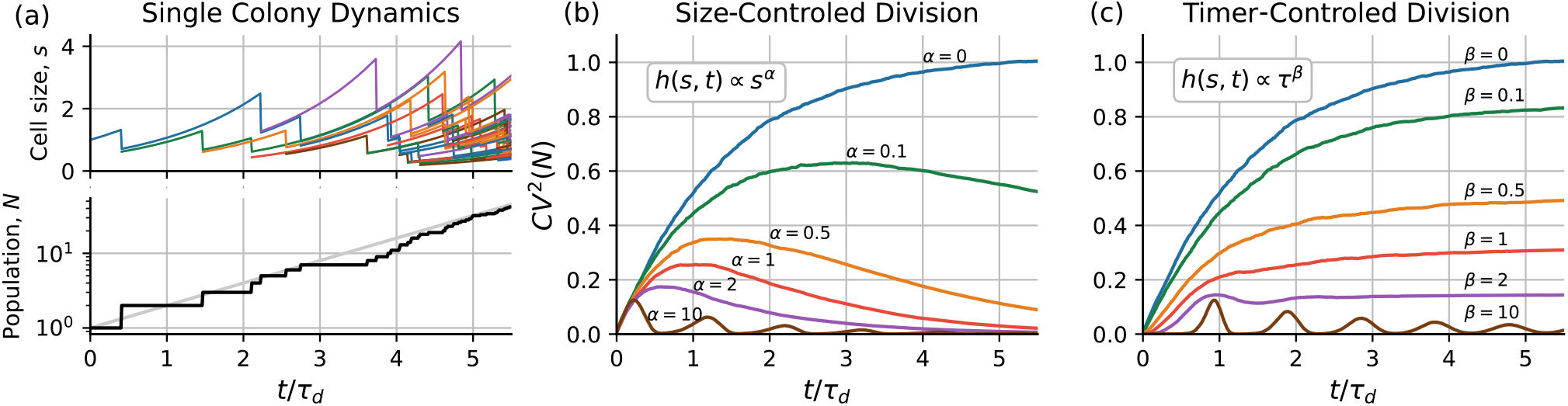
Dynamics of population variability for different division strategies. (a) *Top:* Example of size dynamics among a proliferating population. *Bottom:* Associated stochastic trajectory of the population number *N* over time *t*. (b) Dynamics of population variability *CV* ^2^(*N*) for different division strategies that use a power of size *s*^*α*^ as a reference. (c) *CV* ^2^(*N*) for division strategies that use a power of cell age *τ* ^*β*^ as a reference to decide when to divide. (Parameters: *µ* = ln(2), ⟨*N*⟩|_*t*=0_ = 1, *CV* ^2^(*N*)|*t* = 0 = 0, ⟨*s*⟩|_*t*=0_ = 1, *CV* ^2^(*s*)|_*t*=0_ = 0, *k*, and *γ* were selected as ⟨*τ*⟩ = *τ*_*d*_ = 1)

**TABLE I.** Variables used throughout the article.

| Variable | Interpretation |
| --- | --- |
| $s$ | Cell size. |
| $\mu$ | Growth rate |
| $\tau_d = \frac{\ln(2)}{\mu}$ | Doubling time. |
| $s_b$ | Cell size at the beginning of the process. |
| $\Delta t$ | Time increment between successive algorithm iterations |
| $s_i$ | Cell size in the interval $i$ for the grid in the numerical integrator |
| $\Delta s_i$ | Width of the $i$-th interval |
| $I$ | Total number of $s_i$ intervals |
| $n(s, t)$ | number density function |
| $N(s, t) = n(s, t)ds$ | Individual cells in the interval $(s, s + ds)$. |
| $N(t) = \int n(s, t)ds$ | Cell population at time $t$ |
| $\rho(s, t) = \frac{n(s, t)}{N(t)}$ | Size PDF in the interval $(s, s + ds)$ |
| $x_l(t)$ | Number of cells divided $l$ times at time $t$. |
| $\vec{X} = (x_0, x_1, \dots)$ | Array describing the colony containing all $x_l$. |
| $P(\vec{X})$ | Probability that the colony be in the state $\vec{X}$. |
| $s_l = \frac{s_b e^{\mu t}}{2^l}$ | Cell size after $l$ divisions at a time $t$. |

Next, consider the division timer *τ* as a reference for division, that is, independent of size. In that case, a general model considers the division rate as proportional to the power of the cell age *τ* ^*β*^ with 0 < *β* < ∞:

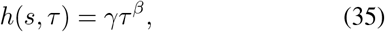

with *γ* being a constant. From this rate, the interdivision times *τ* are distributed following a Weibull distribution [40]:

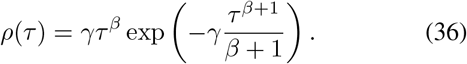

As can be seen in Fig. 6c, *β* = 0 is the constant rate division, and increasing *β* the noise in *N* is more controlled but approaches either a non-null constant or an oscillatory behavior if the control strength (*β*) is high enough. This finite limit shows a fundamental discrepancy with the size-dependent division rate, which goes to zero.

## V. DISCUSSION

Current methods generally model population heterogeneity from two scenarios. In the first, they solve the steady distribution where all moments do not change along time [26], [35]. In the second, the time dynamics can be estimated, but considering a large number of cells. In this way, fluctuations in the cell population are negligible [30]. This article works on the case where these assumptions do not hold: singlecell derived colonies at low population numbers. This limit is essential in applications such as studying how a pathogen invades a host [6], genetic drift in small populations [5], or the survival of small groups to antimicrobial treatment [41].

Here, we analyze the effects of the division strategy on proliferation variability. Studying the transient dynamics of this variability can help us in the discrimination between a size-based division and a timer-based one. Understanding these mechanisms can help to approach phenomenological proliferation models [37] and associate them with molecular mechanisms as in [21], [26]. Furthermore, simply being aware of the differences in the resulting distributions depending on the way the population is followed can prevent error when analyzing population data.

In this work, we present a few details on the additional mechanisms involved in cell division, focusing on the most external variables, such as division rate and splitting noise. However, there are other additional variables, such as the occurrence of multiple events instead of just one to trigger division [17], noise in the growth rate [14], or fluctuations in the initial population [29]. We plan to consider these variables in future publications.

## VI. CONCLUSIONS

This article shows methods for computing the statistical dynamics of both the size and population number in exponentially elongating and proliferating cells. First, we present a fast and efficient way to estimate the size distribution at any time from any starting condition. We observe how different the size distribution is depending on whether we track only one cell after cell division or all the offspring. The distribution considering the first perspective has a higher mean and less variability relative to the second. We modified the algorithm to include stochastic variability in the partitioning position, revealing that the steady distribution has a different shape, although the moment dynamics are essentially the same as the case without stochasticity in partitioning.

This method is valid for a high number of individuals, and therefore it cannot estimate the fluctuations within a colony of a low number of cells. To overcome this, we propose a formalism in which the population is described by an array whose *l*-th component is the number of cells that have *l* divisions. We offer a master equation associated with this system, using numerical methods based on the finite-state projection algorithm to obtain a solution. This article shows how, at low numbers, the size statistics over the colony can differ from those calculated using the PBE.

Finally, we study fluctuations in the population number for colonies of individuals coming from a single ancestor. We observe how if the division is controlled on the basis of the size, the fluctuations in population number go to zero as time passes. Meanwhile, if the division happens with the cell inter-division timer as a reference, these fluctuations tend to be a nonnull constant.

## ACKNOWLEDGMENT

AS is supported by NIH 1R01GM124446-01.

## DATA AVAILABILITY

Codes and data used for this article can be found at https://github.com/canietoa/PopulationFluctuations

## References

[1] J. M. Raser and E. K. O’shea, “Noise in gene expression: origins, consequences, and control,” Science, vol. 309, no. 5743, pp. 2010–2013, 2005.

[2] D. A. Oyarzún, J.-B. Lugagne, and G.-B. V. Stan, “Noise propagation in synthetic gene circuits for metabolic control,” ACS synthetic biology, vol. 4, no. 2, pp. 116–125, 2015.

[3] M. Hashimoto, T. Nozoe, H. Nakaoka, R. Okura, S. Akiyoshi, K. Kaneko, E. Kussell, and Y. Wakamoto, “Noise-driven growth rate gain in clonal cellular populations,” Proceedings of the National Academy of Sciences, vol. 113, no. 12, pp. 3251–3256, 2016.

[4] J. Masel, “Genetic drift,” Current Biology, vol. 21, no. 20, pp. R837–R838, 2011.

[5] D. A. Charlebois and G. Balázsi, “Modeling cell population dynamics,” In Silico Biology, vol. 13, no. 1-2, pp. 21–39, 2019.

[6] C. A. Gilligan and F. van den Bosch, “Epidemiological models for invasion and persistence of pathogens,” Annu. Rev. Phytopathol., vol. 46, pp. 385–418, 2008.

[7] F. Lyu, M. Pan, S. Patil, J.-H. Wang, A. Matin, J. R. Andrews, and S. K. Tang, “Phenotyping antibiotic resistance with single-cell resolution for the detection of heteroresistance,” Sensors and Actuators B: Chemical, vol. 270, pp. 396–404, 2018.

[8] R. Lande, S. Engen, B.-E. Saether, et al., Stochastic population dynamics in ecology and conservation. Oxford University Press on Demand, 2003.

[9] X. Guo, K. P. T. Silva, and J. Q. Boedicker, “Single-cell variability of growth interactions within a two-species bacterial community,” Physical Biology, vol. 16, no. 3, p. 036001, 2019.

[10] S. Abalde-Cela, A. Gould, X. Liu, E. Kazamia, A. G. Smith, and C. Abell, “High-throughput detection of ethanol-producing cyanobacteria in a microdroplet platform,” Journal of The Royal Society Interface, vol. 12, no. 106, p. 20150216, 2015.

[11] D. Volfson, J. Marciniak, W. J. Blake, N. Ostroff, L. S. Tsimring, and J. Hasty, “Origins of extrinsic variability in eukaryotic gene expression,” Nature, vol. 439, no. 7078, pp. 861–864, 2006.

[12] L. Robert, M. Hoffmann, N. Krell, S. Aymerich, J. Robert, and M. Doumic, “Division in escherichia coliis triggered by a size-sensing rather than a timing mechanism,” BMC biology, vol. 12, no. 1, pp. 1–10, 2014.

[13] J. Lin and A. Amir, “The effects of stochasticity at the single-cell level and cell size control on the population growth,” Cell systems, vol. 5, no. 4, pp. 358–367, 2017.

[14] C. Nieto, C. Vargas-Garcia, and J. M. Pedraza, “Continuous rate modeling of bacterial stochastic size dynamics,” Physical Review E, vol. 104, no. 4, p. 044415, 2021.

[15] P.-Y. Ho, J. Lin, and A. Amir, “Modeling cell size regulation: From single-cell-level statistics to molecular mechanisms and population-level effects,” Annual review of biophysics, vol. 47, pp. 251–271, 2018.

[16] S. Iyer-Biswas, C. S. Wright, J. T. Henry, K. Lo, S. Burov, Y. Lin, G. E. Crooks, S. Crosson, A. R. Dinner, and N. F. Scherer, “Scaling laws governing stochastic growth and division of single bacterial cells,” Proceedings of the National Academy of Sciences, vol. 111, no. 45, pp. 15912–15917, 2014.

[17] K. R. Ghusinga, C. A. Vargas-Garcia, and A. Singh, “A mechanistic stochastic framework for regulating bacterial cell division,” Scientific reports, vol. 6, no. 1, pp. 1–9, 2016.

[18] C. Nieto, J. Arias-Castro, C. Sánchez, C. Vargas-García, and J. M. Pedraza, “Unification of cell division control strategies through continuous rate models,” Physical Review E, vol. 101, no. 2, p. 022401, 2020.

[19] C. A. Vargas-Garcia, K. R. Ghusinga, and A. Singh, “Cell size control and gene expression homeostasis in single-cells,” Current opinion in systems biology, vol. 8, pp. 109–116, 2018.

[20] M. Basan, M. Zhu, X. Dai, M. Warren, D. Sévin, Y.-P. Wang, and T. Hwa, “Inflating bacterial cells by increased protein synthesis,” Molecular systems biology, vol. 11, no. 10, p. 836, 2015.

[21] F. Si, G. Le Treut, J. T. Sauls, S. Vadia, P. A. Levin, and S. Jun, “Mechanistic origin of cell-size control and homeostasis in bacteria,” Current Biology, vol. 29, no. 11, pp. 1760–1770, 2019.

[22] P. Wang, L. Robert, J. Pelletier, W. L. Dang, F. Taddei, A. Wright, and S. Jun, “Robust growth of escherichia coli,” Current biology, vol. 20, no. 12, pp. 1099–1103, 2010.

[23] G. Derfel, B. Van Brunt, and G. Wake, “A cell growth model revisited,” Oxford University Research Arhive, 2009.

[24] N. Totis, C. Nieto, A. Küper, C. Vargas-García, A. Singh, and S. Waldherr, “A population-based approach to study the effects of growth and division rates on the dynamics of cell size statistics,” IEEE Control Systems Letters, vol. 5, no. 2, pp. 725–730, 2020.

[25] C. A. Nieto-Acuna, C. A. Vargas-Garcia, A. Singh, and J. M. Pedraza, “Efficient computation of stochastic cell-size transient dynamics,” BMC bioinformatics, vol. 20, no. 23, pp. 1–6, 2019.

[26] C. Jia, A. Singh, and R. Grima, “Cell size distribution of lineage data: analytic results and parameter inference,” Iscience, vol. 24, no. 3, p. 102220, 2021.

[27] J. Lin and A. Amir, “Homeostasis of protein and mrna concentrations in growing cells,” Nature communications, vol. 9, no. 1, pp. 1–11, 2018.

[28] F. F. Mandy, M. Bergeron, and T. Minkus, “Principles of flow cytometry,” Transfusion science, vol. 16, no. 4, pp. 303–314, 1995.

[29] D. Taylor, N. Verdon, P. Lomax, R. J. Allen, and S. Titmuss, “Tracking the stochastic growth of bacterial populations in microfluidic droplets,” Physical biology, 2022.

[30] P. Thomas, “Intrinsic and extrinsic noise of gene expression in lineage trees,” Scientific reports, vol. 9, no. 1, pp. 1–16, 2019.

[31] G. I. Bell and E. C. Anderson, “Cell growth and division: I. a mathematical model with applications to cell volume distributions in mammalian suspension cultures,” Biophysical journal, vol. 7, no. 4, pp. 329–351, 1967.

[32] A. Barizien, M. Suryateja Jammalamadaka, G. Amselem, and C. N. Baroud, “Growing from a few cells: combined effects of initial stochasticity and cell-to-cell variability,” Journal of the Royal Society Interface, vol. 16, no. 153, p. 20180935, 2019.

[33] S. Modi, C. A. Vargas-Garcia, K. R. Ghusinga, and A. Singh, “Analysis of noise mechanisms in cell-size control,” Biophysical journal, vol. 112, no. 11, pp. 2408–2418, 2017.

[34] P. Thomas, “Making sense of snapshot data: ergodic principle for clonal cell populations,” Journal of The Royal Society Interface, vol. 14, no. 136, p. 20170467, 2017.

[35] P. Thomas, “Analysis of cell size homeostasis at the single-cell and population level,” Frontiers in Physics, vol. 6, p. 64, 2018.

[36] B. Munsky and M. Khammash, “The finite state projection algorithm for the solution of the chemical master equation,” The Journal of chemical physics, vol. 124, no. 4, p. 044104, 2006.

[37] B. Houchmandzadeh, “General formulation of luria-delbrück distribution of the number of mutants,” Physical Review E, vol. 92, no. 1, p. 012719, 2015.

[38] K. B. Athreya, P. E. Ney, and P. Ney, Branching processes. Courier Corporation, 2004.

[39] M. Osella, E. Nugent, and M. C. Lagomarsino, “Concerted control of escherichia coli cell division,” Proceedings of the National Academy of Sciences, vol. 111, no. 9, pp. 3431–3435, 2014.

[40] Z. Vahdat, Z. Xu, and A. Singh, “Modeling protein concentrations in cycling cells using stochastic hybrid systems,” IFAC-PapersOnLine, vol. 54, no. 9, pp. 521–526, 2021.

[41] E. Kussell, R. Kishony, N. Q. Balaban, and S. Leibler, “Bacterial persistence: a model of survival in changing environments,” Genetics, vol. 169, no. 4, pp. 1807–1814, 2005.

